# *Trans*-cinnamaldehyde attenuates *Enterococcus faecalis* virulence and inhibits biofilm formation

**DOI:** 10.1101/2021.03.15.435450

**Authors:** Islam A. A. Ali, Jukka P. Matinlinna, Celine M. Lévesque, Prasanna Neelakantan

## Abstract

*Enterococcus faecalis* as an important nosocomial pathogen is critically implicated in the pathogenesis of endocarditis, urinary tract and surgical wound infections. Its major virulence attributes (biofilm formation, production of proteases and hemolytic toxins) enable it to cause extensive host tissue damage. With the alarming increase in enterococcal resistance to antibiotics, novel therapeutics are required to inhibit *E. faecalis* biofilm formation and virulence. *Trans*-cinnamaldehyde (TC), the main phytochemical in cinnamon essential oils has demonstrated promising activity against a wide range of pathogens. Here, we comprehensively investigated the effect of TC on planktonic growth, biofilm formation, proteolytic and hemolytic activities, as well as gene regulation in *E. faecalis*. Our findings revealed that sub-inhibitory concentrations of TC reduced biofilm formation, biofilm exopolysaccharides as well as its proteolytic and hemolytic activities. Mechanistic studies revealed significant down regulation of the quorum sensing *fsr* locus and downstream *gelE*, which are major virulence regulators in *E. faecalis*. Taken together, our study highlights the potential of TC to inhibit *E. faecalis* biofilm formation and its virulence.

## Introduction

The Gram-positive bacterium *Enterococcus faecalis* is a leading cause of endocarditis, urinary tract, surgical wound, and persistent dental root canal infections [1]. Although *E. faecalis* is a commensal of the gastrointestinal tract in humans and animals [2], it is known for its virulence attributes including biofilm formation, production of proteolytic enzymes, and antimicrobial resistance, which complicate the treatment of associated infections [3–5]. The biofilm mode of growth is a survival strategy adopted by microbial cells, thereby securing a higher tolerance to antimicrobials compared to their planktonic counterparts [6, 7]. Slower growth rate of the resident microbes, altered chemical microenvironments, and secretion of a protective extracellular polymeric matrix are responsible for biofilm tolerance to antimicrobials [8]. As a result, higher doses of antimicrobials are required to clear microbes, thus placing a strong selective pressure to develop resistance [9]. Furthermore, sub-inhibitory concentrations of antibiotics such as ampicillin resulted in enhanced *E. faecalis* biofilm development and increased expression of adhesion and biofilm-associated genes [10, 11]. Therefore, interfering with biofilm development during its early stages is considered a promising strategy to avoid the burden of recalcitrant infections.

Regulation of bacterial virulence is mediated by quorum sensing (QS) pathways [12, 13], wherein the population behaviour is synchronized and the expression of regulatory genes is altered in a population density dependent manner [12, 14]. Attenuation of bacterial pathogenicity without affecting bacterial growth is the hallmark of QS quenching as a therapeutic strategy [13]. Therefore, identifying novel anti-QS agents should be proritized to combat biofilm-mediated infections.

Medicinal plants have a vast repertoire of biologically active compounds that are widely used in pharmaceuticals [15, 16]. Such compounds protect plants against pathogens in their natural settings [17]. Interestingly, several phytocompounds have demonstrated anti-QS activity at non-lethal concentrations, and thus can suppress bacterial pathogenicity without triggering resistance phenotypes [18, 19]. *Trans*-cinnamaldehyde (TC) is an aromatic aldehyde, which predominately exists in cinnamon essential oils [20, 21]. It has been shown to have antimicrobial and antibiofilm activities against a wide range of bacterial pathogens including *Pseudomonas aeruginosa* [22, 23], *Escherichia coli* [24, 25], *Staphylococcus aureus* [26,27], *Staphylococcus epidermidis* [28], and *Streptococcus mutans* [29]. There is some evidence to show that TC inhibits growth of *E. faecalis* [30, 31], without inducing an adaptive phenotype [31]. Our pilot studies clearly demonstrated that TC could inhibit these attributes in *E. faecalis* OG1RF strain (ATCC 47077) and ATCC 29212 strains. Therefore, in this study, we performed comprehensive investigations to determine the effects of TC on growth, biofilm formation, biofilm matrix polysaccharides, virulence and gene expression in *E. faecalis*. This study demonstrated the potential of TC to inhibit the virulence attributes of *E. faecalis* including biofilm formation, hemolytic and proteolytic activities at sub-inhibitory concentrations. The anti-virulence effect of TC is possibly mediated via down-regulation of the quorum sensing *fsr* locus and its downstream *gelE* gene.

## Materials and Methods

### Bacterial strains, growth conditions and chemicals

*E. faecalis* reference strains OG1RF (Cyl^-^) and ATCC 29212 (Cyl^+^) were routinely maintained on horse blood agar. All the experiments were performed using the strain OG1RF except the hemolysis assay, which was performed using the strain ATCC 29212, because the latter strain is known to be hemolytic as it harbors the cytolysin operon [32, 33]. Insertion deletion mutants of the *fsr* locus (TX 5241 and TX 5242) of the strain OG1RF were also used [34, 35]. For all experiments, planktonic suspensions were prepared from an overnight culture grown in Brain Heart Infusion (BHI) broth at 37°C. TC and dimethyl sulfoxide (DMSO) were purchased from Sigma Aldrich (St. Louis, MO, USA). All the experiments were performed in triplicates on three independent occasions.

### Planktonic growth assessment

The antimicrobial susceptibility of *E. faecalis* to TC was investigated by determining the minimal inhibitory concentration (MIC) using the broth microdilution assay according to CLSI guidelines [36]. Planktonic suspensions (1 × 10^6^ CFU/ml) were added to microplate wells containing two-fold serially diluted TC (0.059-15 mM) in 0.5% (vol/vol) DMSO. After incubation, the MIC was determined as the lowest concentration of TC that inhibited visible bacterial growth. To investigate the effects of sub-inhibitory (sub-MIC) concentrations of TC on bacterial growth in a time-dependent manner, planktonic cells were incubated with sub-MICs of TC (MIC/8, MIC/4 and MIC/2) in a 96-well polystyrene plate at 37°C. Growth was determined by measuring the optical density (OD 595 nm) of the suspensions every 2 h using a microplate reader (SpectraMax M2, Molecular Devices, LLC, San Jose, CA, USA).

### Biofilm formation assays

Sub-MICs of TC in BHI broth were inoculated with planktonic suspensions in 96-well polystyrene plates and incubated at 37°C for 24 h. After incubation, the planktonic supernatant was carefully removed, and the biofilms were washed three times with sterile phosphate-buffered saline (PBS) to remove the non-adherent cells. The effects of TC on *E. faecalis* biofilm formation was evaluated by quantifying the viable cell counts, metabolic activity, and biomass of developed biofilms.

#### Quantification of viable cell counts

Cell counts were determined using the Colony Forming Units (CFUs) assay. The biofilms were collected by vigorous scraping and pipetting. The collected biofilm suspensions were serially diluted and plated on blood agar. The colonies were counted after incubation at 37°C for 48 h.

#### Assessment of biofilm metabolic activity

To assess the biofilm metabolic activity, a reaction solution of PBS, 1 mg/ml of XTT (2,3-bis (2-methoxy-4-nitro-5-sulfo-phenyl)-2H-tetrazolium-5-carboxanilide) and menadione (70 μg/ml) was freshly prepared at a ratio of 79:20:1. Two hundred microliters of the reaction solution was added to the wells and incubated in dark conditions for 3 h at 37°C [37]. After incubation, 100 μl of the supernatant was transferred to a new plate and the absorbance was measured at 492 nm.

#### Biomass quantification

The biomass of developed biofilms was quantified using the crystal violet assay [38]. Briefly, biofilms were stained with 0.1% (wt/vol) crystal violet (CV) dye for 10-15 min. After staining, the biofilms were washed with sterile PBS to remove the excess dye and left to dry for 15 min at room temperature. The retained dye was solubilized with 95% (vol/vol) ethanol for 10 min, and the absorbance of the supernatant was measured at 570 nm.

### Confocal microscopic imaging of biofilms

Biofilms were developed on plastic coverslips (diameter = 13 mm, thickness = 0.2 mm) (Nunc Thermanox TM, Thermo Fisher Scientific, Waltham, MA, USA) in the presence or absence of sub-MICs of TC at 37°C for 24 h. After incubation, the biofilms were gently washed with sterile PBS to remove the non-adherent cells, and stained using BacLight viability kit TM (Thermo Fisher Scientific) for 30 min. The biofilms were examined using an oil-immersion objective lens (x60) and imaged by confocal laser scanning microscopy (CLSM; Fluoview FV 1000, Olympus, Tokyo, Japan) at five randomly selected points. The biofilm images were analyzed using the Cell-C software to calculate the attached bacterial cells [39].

### Quantification of biofilm exopolysaccharides

Biofilm exopolysaccharide in sub-MIC TC treated biofilms was quantified using the standard phenol sulfuric acid method [40]. Biofilms were allowed to develop in the presence or absence of sub-MICs of TC in a 96 well-polystyrene plate at 37°C for 24 h. After incubation, the planktonic supernatant was carefully removed, and the biofilms were washed with sterile PBS.

Deionized water (20 μl), 5% phenol solution (20 μl) (Sigma Aldrich, St. Louis, MO, USA) and 98% sulfuric acid (100 μl) were added to the wells and incubated at 90°C for 30 min [41]. The absorbance was measured at 492 nm using a microplate reader, and the concentration of biofilm exopolysaccharides was determined using a standard curve generated using glucose standards.

### Extracellular protease production

The activity of extracellular proteases produced by *E. faecalis* was evaluated using the gelatinase plate assay [42]. Briefly, OG1RF planktonic cells (10^7^ CFU/ml) treated with sub-MICs of TC were spot inoculated on agar supplemented with 5% (w/v) gelatin and incubated at 37°C for 24 h. The diameter of opaque zones surrounding the bacterial colonies was measured to determine the effect of TC on extracellular proteases activity.

### Hemolytic activity analysis

The effect of TC on *E. faecalis* hemolytic activity was determined using a previously described protocol with minor modifications [31]. Horse erythrocytes were collected from defibrinated horse blood after centrifugation (500 ×*g*, 10 min) and washed three times with sterile PBS. The planktonic cells of the strain ATCC 29212 (10^7^ CFU/ml) were incubated with or without sub-MICs of TC in presence of 4% (vol/vol) horse erythrocytes at 37°C for 24 h. Our preliminary results showed similar MICs of TC against the strains OG1RF and ATCC 29212. After incubation, control and TC-treated cultures were centrifuged, 100 μl of the supernatant was transferred to a 96-well polystyrene plate before measuring the absorbance at 550 nm using a microplate reader.

### Gene expression analysis of TC-treated biofilms

To study the effects of TC on the expression of quorum sensing-, virulence-, and cell division-associated genes, *E. faecalis* biofilms were allowed to develop in the presence or absence of TC (MIC/8) in a sterile polystyrene plate at 37°C for 24 h. After incubation, the planktonic supernatant was carefully removed, and the biofilms were washed twice with sterile PBS. Biofilms suspensions were centrifuged (9500 *xg*, 10 min) and the collected cells were lysed using lysozyme supplemented with mutanolysin as described previously [43]. Total RNA was extracted using RNeasy Mini Kit (Qiagen, Hilden, Germany) according to the manufacturers’ instructions, and the complementary DNA (cDNA) was prepared from the RNA template using High-Capacity cDNA Reverse Transcription Kit TM (Applied Biosystems, Foster City, CA, USA).

Quantitative Real-Time PCR (qRT-PCR) reactions were run on ABI Step One Plus TM Real-Time PCR System (Applied Biosystems, Foster City, CA, USA). The housekeeping gene 23S rRNA was used as an internal control to which the expression levels of the tested genes were normalized. Relative gene expression was determined by 2^-ΔΔ CT^ method [44]. The sequences of primers used in this study are listed in (S1 Table).

### Statistical analysis

Data from all the phenotypic assays were analyzed using One-way ANOVA and post-hoc Dunnett’s test. Student’s t-test was used to analyze the effect of TC on gene expression. The analyses were performed using IBM SPSS Statistics (Version 26.0, Armonk, NY, USA). P ≤ 0.05 was considered statistically significant.

## Results and Discussion

### Sub-inhibitory concentrations of TC inhibit *E. faecalis* biofilm formation

First, we investigated the antibacterial activity of TC against *E. faecalis* by determining its MIC. The results demonstrated that TC inhibited planktonic growth of the OG1RF strain at 3.7 mM. Ferro et al. demonstrated that TC inhibited the planktonic growth of *E. faecalis* ATCC 19433 and clinical strains at 250 μg/ml (1.88 mM) [31], which is lower than the concentration reported in our study. Difference in the tested strains may explain such a variation. TC has been shown to inhibit the planktonic growth of other bacterial species such as *S. aureus* [31], *Streptococcus pyogenes* [45], and *S. epidermidis* [28], at concentrations close to that observed in this study. The growth kinetics study showed that TC at MIC/2 concentration (1.88 mM), significantly reduced the planktonic growth by at least 32% (p ≤ 0.01, Fig 1), while TC at MIC/4 (0.94 mM) significantly affected the planktonic growth up to 20 h of incubation (p ≤ 0.05, Fig 1). By contrast, the MIC/8 concentration (0.47 mM) had no significant effect on planktonic growth (p > 0.05, Fig 1). The estimated generation time was 61 min for the control planktonic culture, while it was 55, 63 and 100 min for TC-treated cultures at 0.47, 0.94 and 1.88 mM respectively.

**Fig 1.**
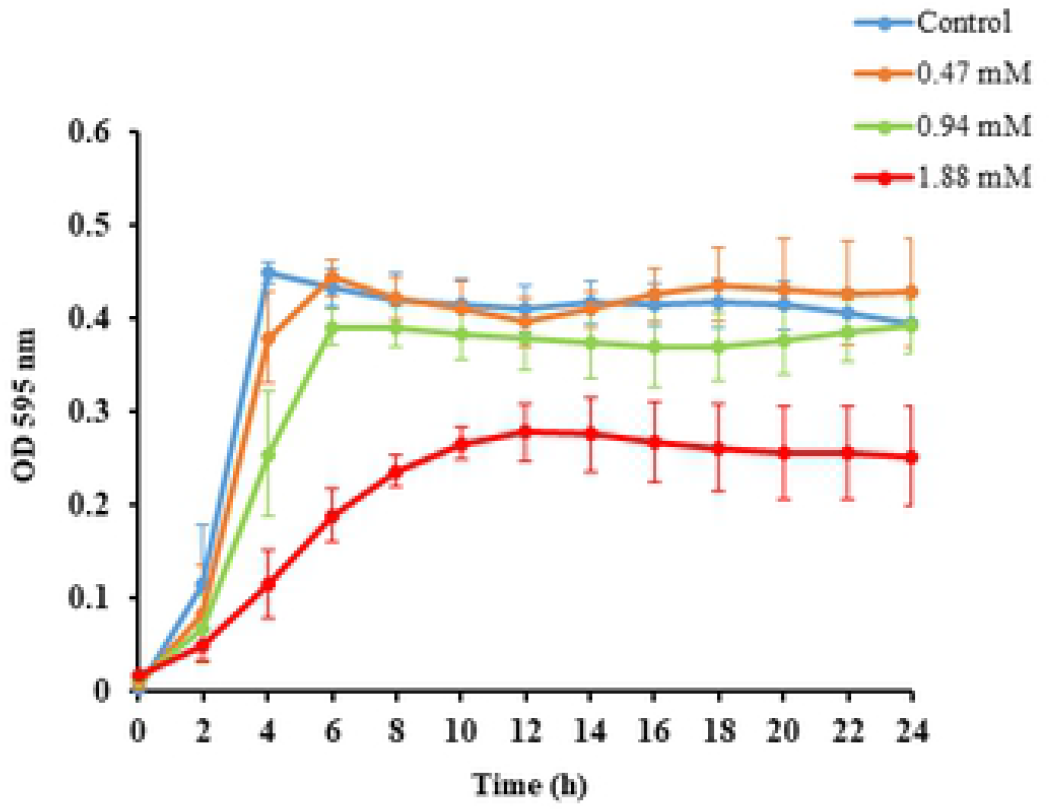
Effect of sub-MICs of TC on planktonic growth kinetics of *E. faecalis*. The growth kinetics plot shows that TC at 0.47 mM had no significant effect on planktonic growth over the 24 h period, compared to untreated controls (p > 0.05).

We then investigated the effect of TC on *E. faecalis* biofilm formation by evaluating the viable cell counts, metabolic activity and biomass of biofilms developed in the presence or absence of sub-MICs of TC. The results showed a significant dose dependent reduction in log_10_

CFU/ml of biofilm cells (p ≤ 0.05, Fig 2a). Ferro et al. demonstrated that TC at MIC/2 and MIC/4 decreased the number of *E. faecalis* cells attached to a catheter model [31]. A similar pattern was observed in the XTT assay, which showed a reduction in the metabolic activity by 12%, 31% and 51% in biofilms developed at 0.47, 0.94 and 1.88 mM respectively. This reduction was significant at 0.94 and 1.88 mM (p ≤ 0.01, Fig 2b), and insignificant at 0.47 mM (p > 0.05, Fig 2b). These findings indicate that 0.47 mM TC potentially interferes with the adhesion of planktonic cells, thus decreasing the number of biofilm cell counts as shown by the CFUs assay, without affecting their viability as demonstrated by the metabolic activity assay.

**Fig 2.**
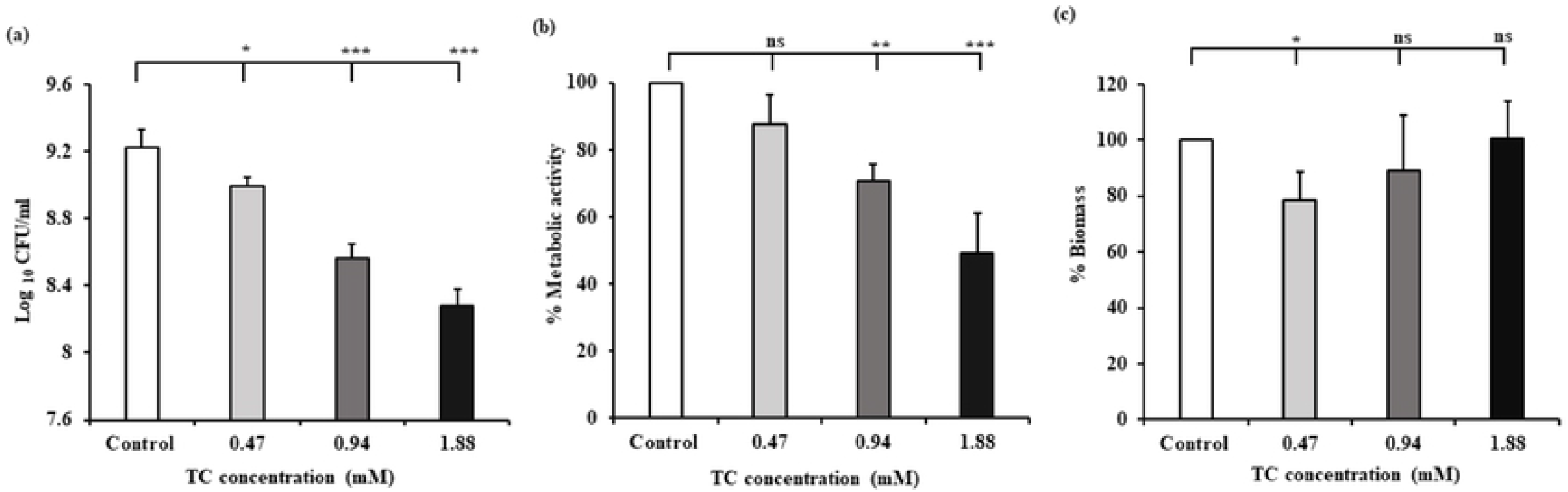
(a) Cell count, (b) metabolic activity, and (c) biomass of *E. faecalis* biofilms developed at sub-MICs of TC. Cell count is expressed as log10 CFU/ml, while the metabolic activity and biomass are expressed relative to the control (100%). * denotes p ≤ 0.05, ** denotes p ≤ 0.01, *** denotes p ≤ 0.001, ns denotes non-significant difference compared to control (p > 0.05).

Agreement in the results of XTT and CFUs assay was observed at 0.94 and 1.88 mM, while such agreement was not observed at 0.47 mM. Variation between the results of XTT and CFU assays has been addressed previously, wherein the XTT absorbance showed a linear response when the logarithm of cell numbers ranges from 4.5-7.5 [46]. Furthermore, a substantial increase in XTT absorbance has been reported above 10^8^ cells/ml compared to a steady increase in XTT absorbance at lower cell number [47]. Thus, a clear-cut linear relation between XTT absorbance and cell numbers cannot be necessarily assumed, and this methodological issue should be taken into consideration when such studies are performed.

We then evaluated the biomass of *E. faecalis* biofilms developed at sub-MICs of TC using the CV assay. A concentration of 0.47 mM significantly reduced the biomass compared to control (p ≤ 0.01, Fig 2c). The biomass of biofilms developed at 0.94 and 1.88 mM were not significantly different from the control (p > 0.05, Fig 2c), which agree with the results reported by Ferro et al. [31], wherein TC had no effect on biofilm formation by clinical *E. faecalis* isolates. Despite the reduction in the number of viable biofilm cells at 1.88 mM observed in our study, no corresponding reduction in the biomass was observed. This is similar to previously reported findings, wherein MIC/2 of TC significantly reduced the number of *S. aureus* viable biofilm cells without being able to reduce the overall biomass [31]. Instead, the biomass was increased probably due to accumulation of dead cells.

Reduction of the biofilm biomass by TC at 0.47 mM as observed in our study corroborate with previous studies, wherein sub-MICs of TC reduced biofilm formation by *S. mutans* [29], *S. pyogenes* [48], and *P. aeruginosa* [22, 23, 49]. In some of these studies, reduction in biofilm formation was associated with suppression of QS pathways in the relevant microorganisms [29, 49]. Therefore, we hypothesize that reduction of *E. faecalis* biofilm formation by TC at 0.47 mM could be mediated via suppression of QS systems. *E. faecalis* has three main QS systems - Fsr, Cytolysin and LuxS [50]. The Fsr QS system contributes to biofilm formation via production of gelatinase [51]. Hence, we tested the effect of TC at 0.47 mM on biofilm formation using two *fsr* mutants TX 5241 and TX 5242, which are characterized by insertion deletion of *fsrB* and *fsrC* genes respectively [34, 35]. The results showed that TC was unable to reduce biofilm formation by the *fsr* mutants (p > 0.05, Fig 3), thus demonstrating that the Fsr QS system is targeted by the biofilm inhibitory concentration of TC and the effect of TC on *E. faecalis* biofilm formation is not evident when the *fsr* QS genes are inactivated.

**Fig 3.**
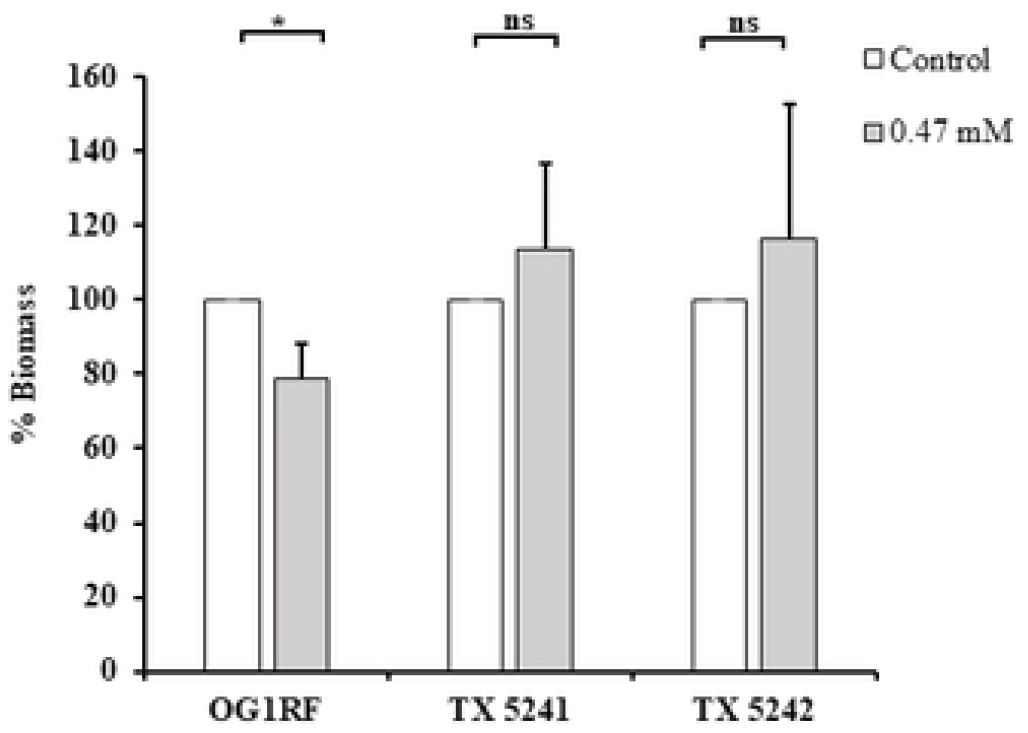
Effect of TC (0.47 mM) on biofilm formation by the OG1RF and *fsr* mutants (Insertion deletion *fsrB* and *fsrC* mutants, TX 5241 and TX 5242 respectively). In each tested strain, the biomass developed in presence of TC was normalized to its respective control (100%). * denotes p ≤ 0.05, ns denotes non-significant difference compared to control (p > 0.05).

To elucidate the antibiofilm effect of TC on *E. faecalis*, we investigated the overall structure of *E. faecalis* biofilms using CLSM. The mean attached bacterial cells was significantly reduced from 13 x 10^3^ cells/mm^2^ in control biofilms to 2 – 3.6 x 10^3^ cells/ mm^2^ in biofilms developed at sub-MICs of TC (p ≤ 0.001, Fig 4b), leaving considerable areas of the substrate devoid of biofilm (Fig 4a). Similar results have been demonstrated for *S. mutans* and *S. pyogenes* biofilms developed at sub-inhibitory concentrations of TC [29, 48].

**Fig 4.**
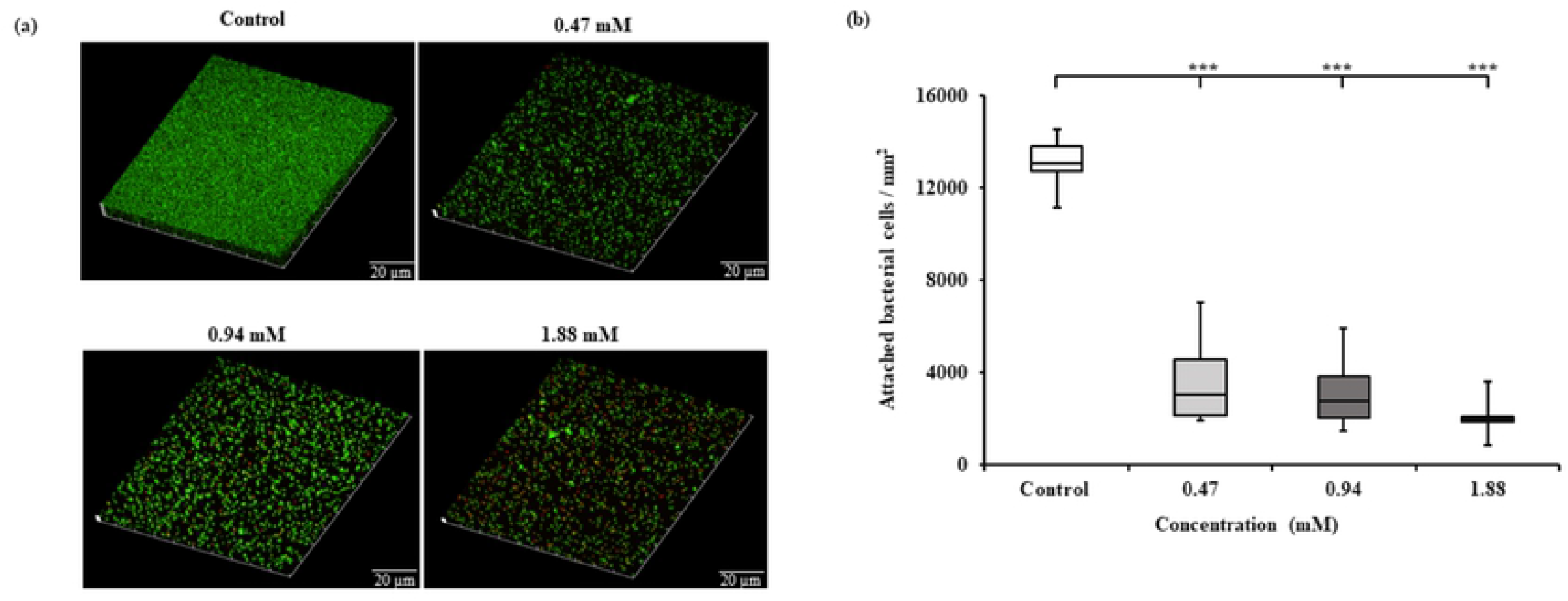
CLSM investigation of *E. faecalis* biofilms developed in presence or absence of TC. **(a)** Three-dimensionally reconstructed images showing reduction of *E. faecalis* biofilm formation by TC. At sub-MICs of TC, the biofilm structure was disrupted and demonstrated reduced density compared to intact and dense control biofilms. **(b)** Sub-MICs of TC significantly reduced the total number of attached bacterial cells/mm^2^, compared to control indicating the inhibition of microbial adhesion and biofilm formation. *** denotes p ≤ 0.001.

### Sub-inhibitory concentration of TC reduces *E. faecalis* biofilm exopolysaccharides

Biofilms are structurally complex communities, in which the cells are embedded in an extracellular polymeric substance (EPS), which contains polysaccharides, proteins, and nucleic acids as its major constituents [52]. Polysaccharides contribute to bacterial adhesion, aggregation, biofilm stability, nutrients acquisition and protection against antimicrobials [53]. Therefore, we investigated the effect of sub-MICs of TC on exopolysaccharides content of *E. faecalis* biofilms. The results demonstrated a significant reduction in the exopolysaccharides of biofilms developed at 0.47 mM of TC (75 ± 15 μg/ml) compared to control biofilms (107 ± 28 μg/ml) (p ≤ 0.01). Such a reduction in exopolysaccharides provides a plausible explanation of the reduced biomass at 0.47 mM as shown by the CV assay. The results of exopolysaccharides and metabolic activity assays in our study support the previously reported findings that TC inhibited biofilm formation by reducing exopolysaccharides production, without affecting the viability of biofilm cells [54]. TC reduced exopolysaccharides production by *Listeria monocytogenes* in accordance with the results of our study [55]. In our study, the biofilm exopolysaccharides at 0.94 (119 ± 21 μg/ml) and 1.88 mM (136 ± 38 μg/ml) were not significantly different from control biofilms (p ≥ 0.05), which explain the insignificant difference in biomass at 0.94 and 1.88 mM compared to control biofilms as shown by CV assay (Fig 2c).

### TC reduces the proteolytic and hemolytic activities of *E. faecalis*

Extracellular proteases are putative virulence factors, which play an important role in the pathogenicity of *E. faecalis* [56]. Gelatinase and serine protease contribute to *E. faecalis* biofilm formation [51], and degradation of host tissues [57, 58]. Thus, we asked if sub-MICs of TC could affect the activity of such proteases. The results showed that TC at 1.88 mM significantly decreased the diameter of proteolytic zones surrounding the bacterial colonies (22.3 ± 0.9 mm in TC group vs 24.5 ± 0.8 mm in control group, p ≤ 0.05). It has been shown that TC reduced protease activity of *S. pyogenes* [48].

Cytolysin (or hemolysin) is an exotoxin, which lyses human erythrocytes, intestinal epithelial and retinal cells, thereby implicated in the pathogenesis of enterococcal infections [59]. The widespread prevalence of hemolysis phenotype in clinical isolates emphasize cytolysin as a promising target for future development of anti-infective agents [60, 61]. Therefore, we investigated the effect of TC on the hemolytic activity of *E. faecalis*. The results showed that TC reduced the hemolytic activity of *E. faecalis* by 21%, 39% and 64% at 0.47, 0.94 and 1.88 mM respectively compared to control (p ≤ 0.001). This corroborates with a previous finding that sub-MIC of TC reduced the hemolytic activity of a clinical isolate of *E. faecalis* [31]. It is interesting to note that sub-MICs of TC also reduced hemolysis induced by *S. aureus* [39], and *P. aeruginosa* [23], indicating that TC has potential to inhibit virulence in a wide range of Gram-positive and Gram-negative pathogens.

### TC suppresses the Fsr quorum sensing pathway and the downstream *gelE* gene

Based on the results of the observed phenotypes, sub-inhibitory concentrations of TC reduced *E. faecalis* biofilm formation, altered the structure of developed biofilms, and attenuated its proteolytic and hemolytic activities. Therefore, in order to understand the underlying mechanisms, we investigated the effect of TC on the expression of quorum sensing-, virulence-, and cell division-associated genes using qRT-PCR. The interest in the *fsr* locus arises from previous findings which showed that the Fsr system was the only two-component signal transduction system in *E. faecalis*, which affects biofilm formation when inactivated [51, 62]. In addition, the Fsr system has an important role in the regulation of surface proteins and several metabolic pathways in *E. faecalis* [63]. The presence of *fsr* locus in 100% of endocarditis isolates and over 50% of fecal isolates strongly suggest that this system is a promising target to develop non-toxic inhibitors, thus attenuating the pathogenicity of *E. faecalis* [64].

The *fsrB* gene encodes a trans-membrane protein that process a propeptide to generate a peptide pheromone, while *fsrC* gene encodes a histidine kinase sensor that responds to the peptide-signaling molecule, phosphorylates its response regulator, and subsequently activates the transcription of the *gelE-sprE* operon [65]. The Fsr QS system contributes to biofilm formation and virulence of *E. faecalis* via gelatinase production [51, 66]. Our results showed that TC at 0.47 mM significantly downregulated the expression of *fsrB* and *fsrC* compared to the control (p ≤ 0.001, Fig 5). TC has been demonstrated to down-regulate the expression of quorum sensing genes in other Gram-positive bacteria such as *S. mutans* [29, 67], and *L. monocytogenes* [55], indicating that TC has potential to inhibit virulence in different species through multiple pathways.

**Fig 5.**
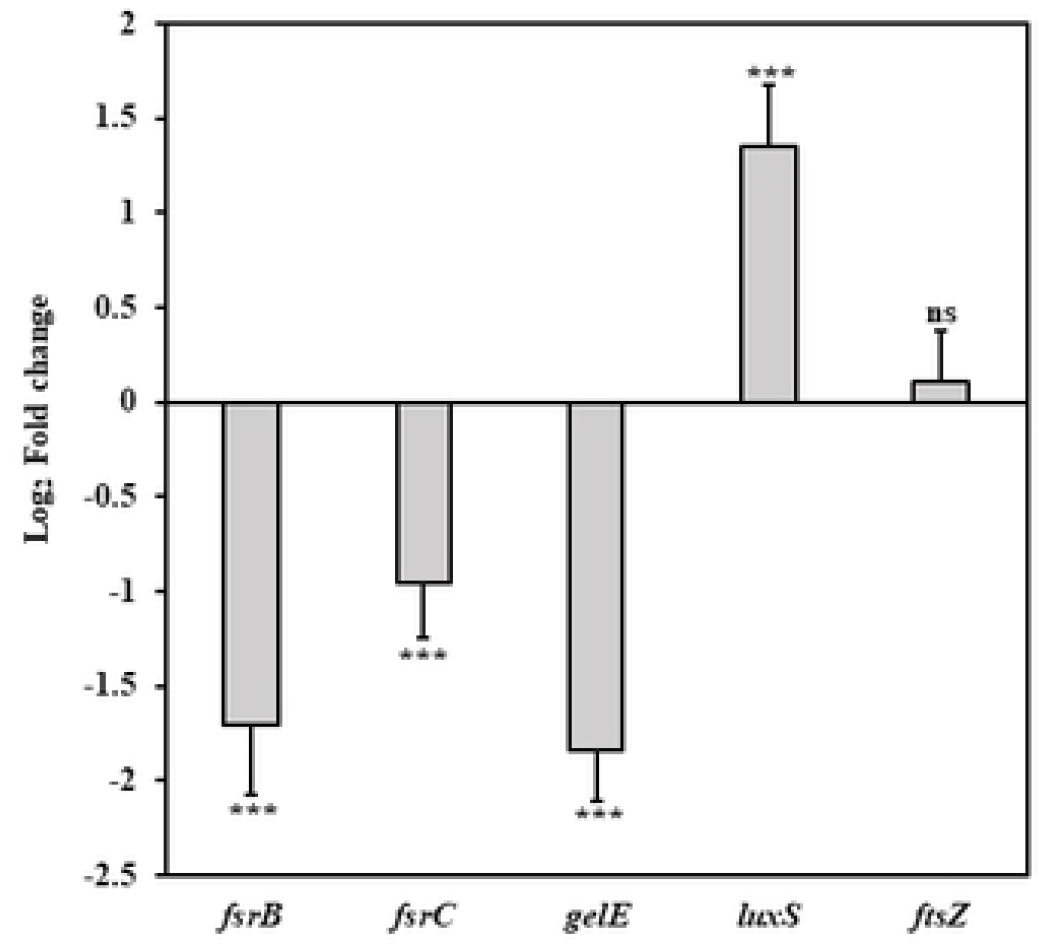
Effect of sub-MIC of TC (0.47 mM) on gene expression in *E. faecalis*. *** denotes p ≤ 0.001, ns denotes no significant difference compared to control (p > 0.05).

Downstream to the *fsr* locus, *gelE* regulates the production of gelatinase, an extracellular metalloprotease, which contributes to biofilm formation via fratricide mediated cell lysis and release of extracellular DNA [68]. The importance of *gelE* for *E. faecalis* pathogenicity was emphasized by its widespread prevalence in isolates from food and clinical sources [69, 70]. In our study, we observed that the transcription of *gelE* was significantly downregulated at the concentration 0.47 mM compared to the control (p ≤ 0.001, Fig 5). In accordance with our results, TC has been shown to reduce the transcription of *speB* and *lasA* genes that encode proteases production in *S. pyogenes* and *P. aeruginosa* respectively [48, 49]. The results of the gene expression analysis corraborate the results of phenotypic biofilm formation assays done on the wild-type OG1RF strain (Fig 2c) and *fsr* mutants (Fig 3). The findings from these assays collectively support the hypothesis that suppression of the *fsr* locus and its downstream *gelE* is a possible mechanism to elucidate the inhibitory effect of TC on biofilm formation in *E. faecalis*.

The LuxS/Autoinducer-2 (AI-2) dependent QS system is a universal signaling pathway involved in interspecies communication, regulation of virulence factors and host-microbe interactions [12, 71]. In this system, *luxS* encodes a metalloenzyme involved in the production of AI-2 [72]. The results in our study showed that a sub-MIC of TC significantly upregulated the expression of *luxS* (p ≤ 0.001, Fig 5). By contrast, expression of *luxS* in *S. pyogenes* and *S. mutans* was downregulated by TC [48, 67]. TC did not affect the AI-2 production but reduced the bioluminescence in the gram-negative *Vibrio harveyi* by targeting the DNA binding activity of the transcriptional regulator LuxR [54, 73]. These contradictory results can be explained by differences in the quorum sensing-regulated phenotypes between bacterial species, and even between the strains of the same species [74]. Such strain-related variations have been reported regarding the role of *luxS* in *E. faecalis*. While the deletion of *luxS* in the strain ATCC 33186 resulted in increased biofilm formation [75], it resulted in opposite effects in the strain ATCC 29212 [76], and this reduction was only significant in 48h-old biofilms.

Inhibition of cell division is one of the proposed antibacterial mechanisms of action of TC [77, 78]. Therefore, we studied the effect of TC on the expression of *ftsZ*, which encodes a key cytoplasmic protein in cell division. Our results showed that the sub-MIC of TC did not significantly affect the expression of *ftsZ* (p > 0.05, Fig 5). TC has been shown to inhibit bacterial cell division by binding to the C-terminal region of FtsZ, thus perturbing the morphology of Z-ring, and interfering with its assembly dynamics [79].

To the best knowledge of the authors, this is the first study to provide comprehensive evidence on the concentration-dependent effects of TC on biofilm inhibition, reduction of biofilm exopolysaccharides and virulence inhibition in *E. faecalis*. However, we investigated the effect of TC on biofilm formation and virulence factors only in the *fsr* positive strain OG1RF. Future investigations should be performed on several clinical isolates with different quorum sensing and virulence profiles. Transcription of genes encoding surface adhesins and other virulence determinants after treatment with TC should also be studied in future work.

## Conclusions

The results of this study highlighted that sub-inhibitory concentrations of *trans*-cinnamaldehyde reduced *E. faecalis* biofilm formation, biofilm exopolysaccharides, altered the structure of developed biofilms, and attenuated the proteolytic and hemolytic activities without inhibiting the bacterial growth. TC also significantly downregulated genes related to the Fsr quorum sensing system in *E. faecalis*.

## Acknowledgments

Islam A.A. Ali is supported by the postgraduate fellowship from the University of Hong Kong, and this study is a part of his doctoral thesis. The authors gratefully acknowledge Professor Barbara E. Murray (Division of Infectious Diseases, Department of Medicine, University of Texas Medical School, Huston, Texas, USA) for gifting us the *fsr* mutant strains. The authors sincerely thank Ms. Becky Cheung, Central Research Laboratories, Faculty of Dentistry, University of Hong Kong for the technical expertise with the confocal microscopic experiment.

## Author Contributions

**Conceptualization:** Prasanna Neelakantan, Celine M. Lévesque, Islam A. A. Ali.

**Methodology:** Prasanna Neelakantan, Islam A. A. Ali.

**Investigation:** Islam A. A. Ali.

**Data curation:** Islam A. A. Ali.

**Resources:** Prasanna Neelakantan, Jukka P. Matinlinna.

**Writing - original draft preparation:** Islam A. A. Ali, Prasanna Neelakantan.

**Writing - review and editing:** Islam A. A. Ali, Prasanna Neelakantan, Celine M. Lévesque, Jukka P. Matinlinna.

**Supervision:** Prasanna Neelakantan, Celine M. Lévesque, Jukka P. Matinlinna.

**Project Administration:** Prasanna Neelakantan.

## Supporting Information

**S1 Table. Sequence of primers used in the qRT-PCR analysis.**

